# How reliable are estimates of key parameters in viral dynamic models?

**DOI:** 10.1101/2023.08.17.553792

**Authors:** Carolin Zitzmann, Ruian Ke, Ruy M. Ribeiro, Alan S. Perelson

## Abstract

Mathematical models of viral infection have been developed and fit to data to gain insight into disease pathogenesis for a number of agents including HIV, hepatitis C and B virus. However, for acute infections such as influenza and SARS-CoV-2, as well as for infections such as hepatitis C and B that can be acute or progress to being chronic, viral load data are often collected after symptoms develop, usually around or after the peak viral load. Consequently, we frequently lack data in the exponential phase of viral growth, i.e., when most transmission events occur. Missing data may make estimation of the time of infection, the infectious period, and parameters in viral dynamic models, such as the cell infection rate, difficult. Here, we evaluated the reliability of estimates of key model parameters when viral load data prior to the viral load peak is missing. We estimated the time from infection to peak viral load by fitting non-linear mixed models to a dataset with frequent viral RNA measurements, including pre-peak. We quantified the reliability of estimated infection times, key model parameters, and the time to peak viral load. Although estimates of the time of infection are sensitive to the quality and amount of available data, other parameters important in understanding disease pathogenesis, such as the loss rate of infected cells, are less sensitive. We find a lack of data in the exponential growth phase underestimates the time to peak viral load by several days leading to a shorter predicted exponential growth phase. On the other hand, having an idea of the time of infection and fixing it, results in relatively good estimates of dynamical parameters even in the absence of early data.

## Introduction

In a typical acute infection, the viral load initially increases exponentially, reaches a peak, and then declines. The same pattern is seen in infections, such as hepatitis C, that can progress from acute to chronic, where the decline does not necessarily lead to elimination of the virus. The viral load frequently correlates with a person’s infectiousness and thus the probability of viral transmission [1–6]. Understanding the viral dynamics throughout the course of infection, including prior to the viral peak, is critical to understanding viral transmission [7–9]. However, viral load data is often obtained from settings where infected individuals are tested and identified days after symptoms develop [3,10–12]. In the case of SARS-CoV-2, symptom onset is usually around the peak viral load and corresponds to the time when a person is highly contagious [1]. This leads to a lack of data collected during the exponential growth phase. Additionally, the exact time of pathogen exposure is often unknown, or estimates of infection times are often based on incomplete data.

Many previous models were fit to data from observational studies with missing data prior to the peak viral load and thus mostly with unknown times of infection, which may lead to uncertainties in estimates of the incubation period and, as we show below, in estimates of viral dynamic model parameters. Here, we present a mathematical analysis estimating key parameters of viral dynamic models from a data set from the National Basketball Association (NBA) where testing for SARS-CoV-2 was done on a regular basis irrespective of infection status, and including pre-peak viral load assessment [10,11]. The time peak viral load was defined as *t*=0 in this dataset [10,11] and thus the time of infection, *t*_inf_, is negative and denotes the number of days before the viral peak that infection was estimated to have taken place.

We found that the cell infection rate and virus production rate are crucial parameters in viral dynamic models needed to reliably estimate the dynamics of the exponential growth phase. If data is missing in the first few days post infection, knowing both parameters led to similar predictions of viral load as having frequent viral load measurements in the exponential growth phase. Alternatively, knowledge of the time of infection (e.g., from epidemiological evidence) or assuming a given duration until peak viral load is attained (e.g., in SARS-CoV-2 assuming the median of 5 days [13,14]) represent good alternatives to estimating infection times and yields consistent population parameter estimates.

## Methods

### Mathematical Model

A mathematical model often used to study acute infections is the target cell limited model (TCLM) with an eclipse phase, which was introduced to study influenza infection [15]. This model has been used to study various acute infections, such as Zika, dengue, influenza A, West Nile virus, Ebola, and SARS-CoV-2 [1,15–19], due to its simplicity and the small number of parameters.

The TCLM describes the dynamics of target cells, i.e., cells susceptible to infection, *T*, infected cells in the eclipse phase that are not yet virus-producing, *E*, virus-producing infected cells, *I*, and virus, *V*. The TCLM has also been augmented by including an innate immune response that has provided a better description of influenza and SARS-CoV-2 infection dynamics [1,15]. In this model, we include a population of cells that are refractory to infection, which for simplicity, we call refractory cells, *R*, and call the model the refractory cell model (RCM). Refractory cells are in an antiviral state induced by the innate immune response mediated by type I and type III interferons [20–22]. The following system of ODEs gives the dynamics of the five populations of the RCM:

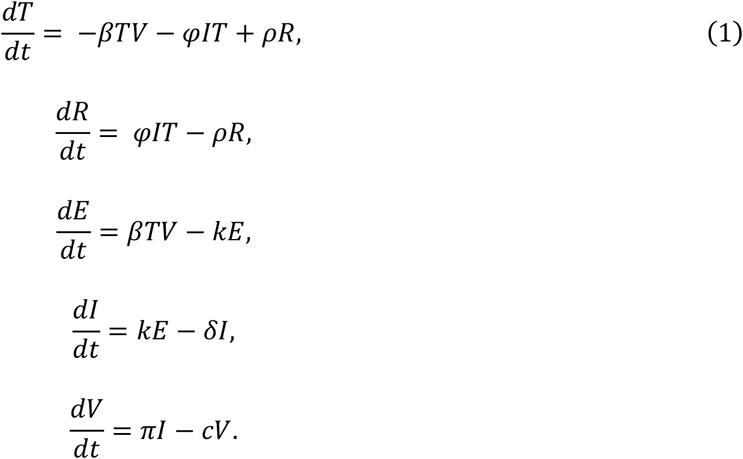

The TCLM is similar and to obtain it we just remove the *dR*/*dt* equation, and the terms *φIT* and *ρR* in the *dT/dt* equation (see equation S1 in S1 Text). In the model, target cells, *T*, become infected by virus with rate constant *β* and then enter the eclipse phase, *E*, which lasts for an average duration 1/*K* during which time they produce no virus. At the end of the eclipse phase cells become productively infected cells, *I*, produce virus with rate constant *π* and die with rate constant *δ*. Note that the average infected cell lifespan is 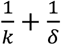. Finally, virus, *V*, is cleared with first-order rate constant *c* (Fig 1).

**Figure 1:**
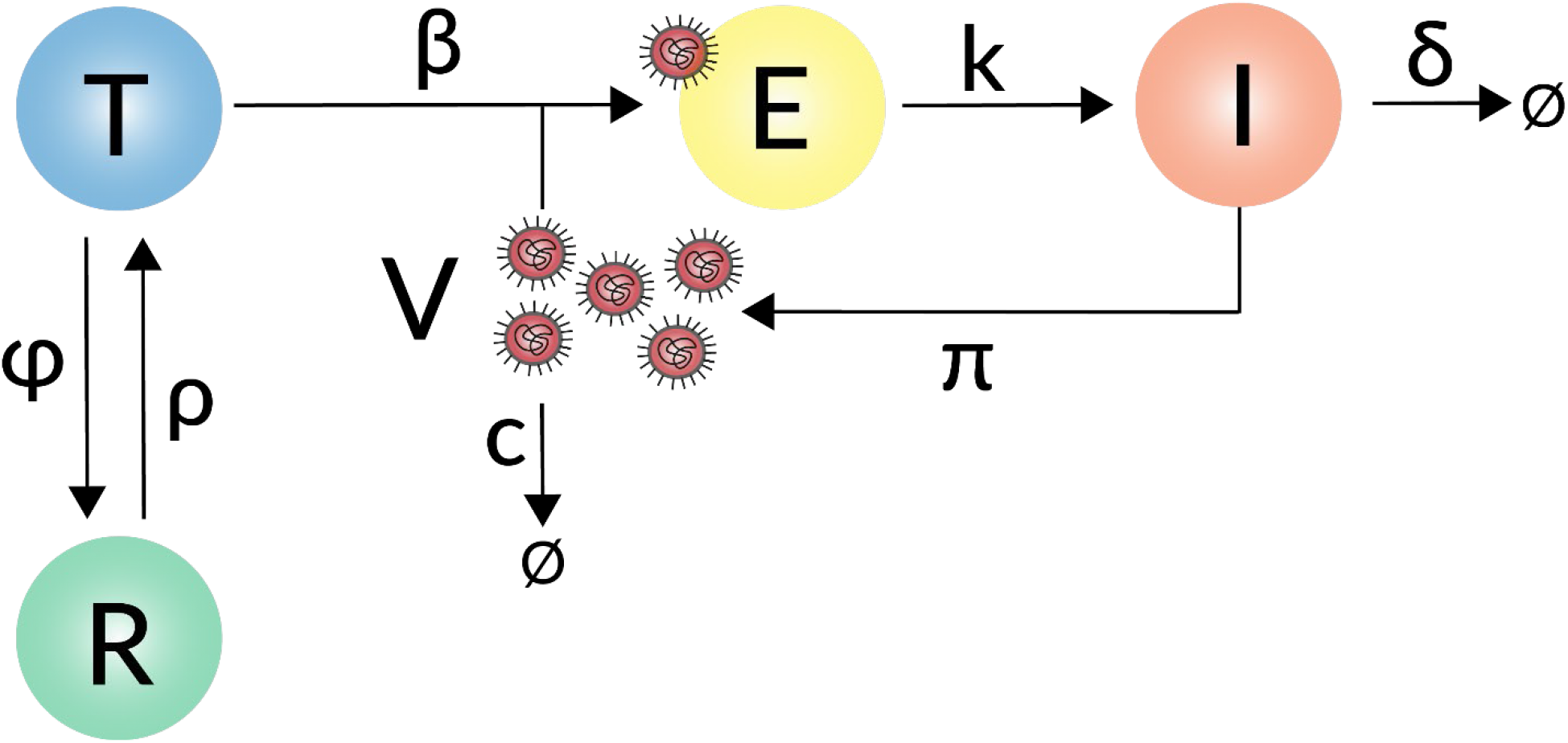
Schematic illustration of the refractory cell model. A susceptible target cell, T, is infected by virus, V, with the infection rate constant β. Infected cells in the eclipse phase, E, become actively virus producing cells, I, with the transition rate constant K. I produce virus with production rate constant π or die with degradation rate δ. Virus is cleared with clearance rate cc. In the refractory cell model, in addition we also account for the innate immune response, which turns susceptible cells into refractory cells, R, with constant rate φ, which are in an antiviral state and refractory to infection. However, refractory cells can become susceptible to infection with constant rate ρ.

Infection induces the release of interferons that may establish an antiviral state in non-infected target cells. For simplicity, we do not explicitly include interferons but model their effect as proportional to the number of infected cells. With per capita rate *φI*, target cells enter the refractory state and leave it with first-order rate constant *ρ*, making them again susceptible to infection (Fig 1).

Consistent with our previous work [23], we fixed the initial target cell population at the time of infection (*t*_*inf*_) to *T*(*t*_*inf*_)= 8 x 10^7^ cells and assumed the initial refractory cell population *R(t*_*inf*_) = 0. The virus concentration that initiates infection is hard to estimate. Thus, as has been done previously [24] we set *V*(*t*_*inf*_) = 0 and start the infection with one infected cell in the eclipse phase: *E*(*t*_*inf*_) = 1 [23], which was the value that led to the best model fit according to our sensitivity analysis (S1 Table). *In vitro* experiments have shown that it usually takes 4–8 hours before an infected cell starts to produce SARS-CoV-2 [15,25], yielding a rate of transition out of the eclipse phase of *K* = 4 *d*^−1^. Further, we fixed the virus clearance rate *c* = 10 *d*^−1^ [25].

For the above models, the basic reproductive number (*R*_0_) is given by

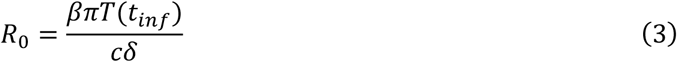

which corresponds to the average number of cells infected by one single infected cell at the start of infection.

For model comparisons below, we calculate the root-mean-square error (RMSE)

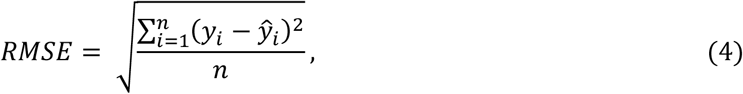

where *y*_*i*_ are the actual measurements and 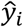 are model predictions at time *i* for all *n* non-censored data points, i.e., viral load measurements above the detection limit (LOD).

### Parameter estimation, model selection, and model analysis

Fitting the RCM to data was implemented using the non-linear mixed effects modeling framework in *Monolix* (lixoft.com) and *R* (r-project.org) using Monolix’s R-functions. We conducted 100 different parameter estimation rounds with randomly chosen initial parameter values uniformly distributed within the following ranges: *t*_*inf*_ ∈ [–8, –5] days, *β* ∈ [10^−8^, 10^−5^] mL/RNA copies/days, *δ* ∈ [0.1, 3] 1/day, *π* ∈ [1, 100] RNA copies/mL/day, *φ* ∈ [10^−8^, 10^−4^] 1/cell/day, and *ρ* ∈ [10^−3^, 10^−1^] 1/ day. The best model fit was selected from the 100 different rounds of parameter estimation by comparing the negative log-likelihood (-LL) and the RMSEs. Note that the randomly chosen initial parameter values serve the purpose of covering a larger parameter search space. However, the estimated parameter values are not necessarily in the defined ranges.

### Data and data collection scenarios

We used published data from the National Basketball Association (NBA) to estimate model parameters, where unvaccinated individuals were regularly tested during the NBA tournament in 2020 and 2021 [10,11]. We selected 25 unvaccinated individuals from this cohort with frequent viral load measurements, i.e., individuals with four or more viral load measurements above the limit of detection (LOD), representing the entire course of infection (viral load up-slope, peak, and down-slope). On average (± standard deviation), the 25 selected individuals had 9.8 ± 3.8 viral load measurements above the LOD with 3.1 ± 1.6 data points obtained during the up-slope, one measurement representing the observed peak viral load, and 5.7 ± 3.0 measurements obtained during the post-peak down-slope. We used this *“*entire course of infection*”* data set to estimate the median time to the measured peak viral load and to study the dynamics of the acute infection (S1 Fig).

## Results

### SARS-CoV-2 dynamics: The course of infection

The RCM fits the entire course of infection of the 25 selected individuals and describes both the initial exponential viral growth and subsequent virus clearance (Fig 2).

**Figure 2:**
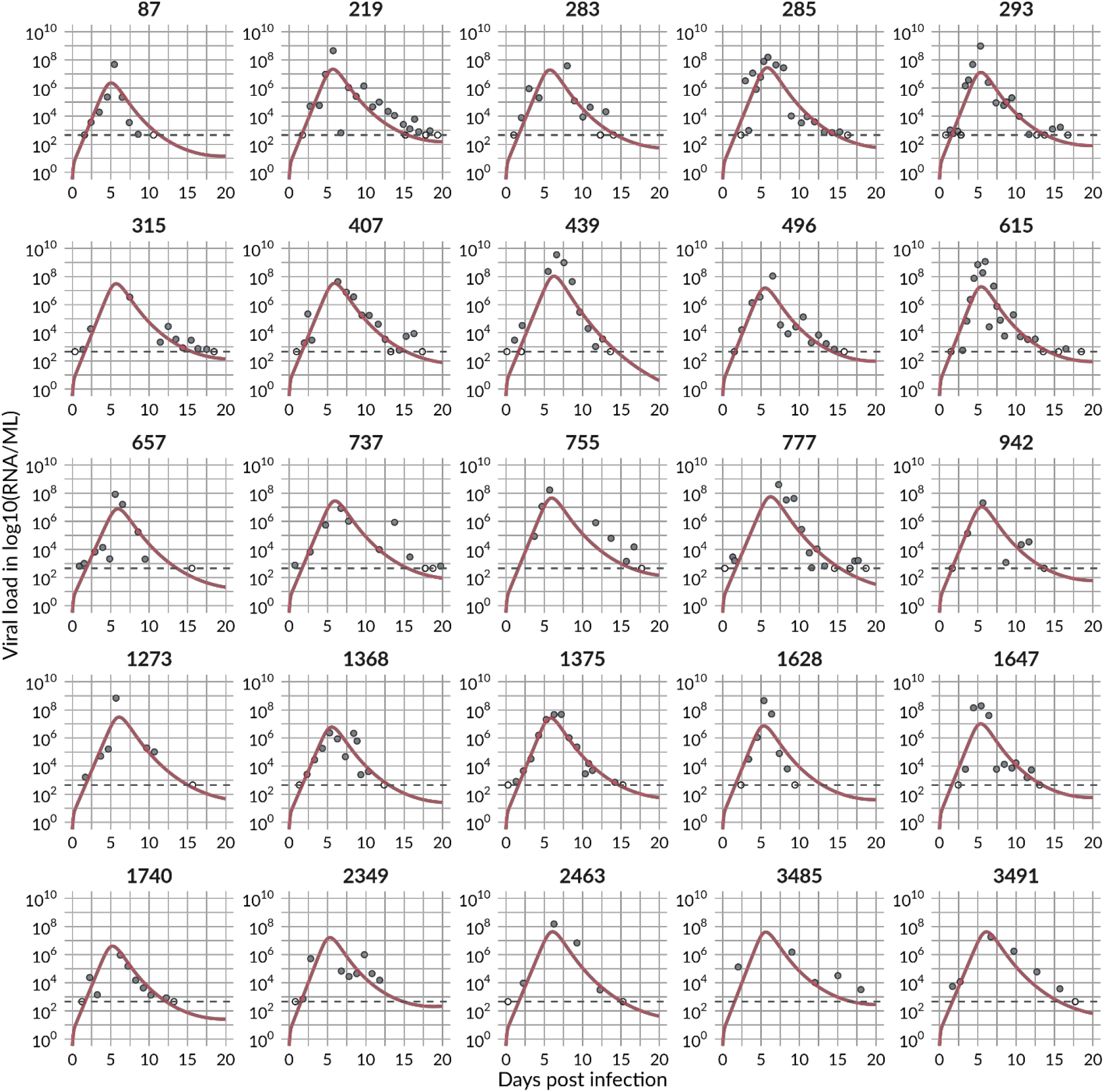
The best RCM fit to viral load measurements of 25 selected individuals. Filled circles are 138 measurement points, and non-filled circles are censored and below the detection limit (dotted grey line).

We estimated the time of infection at a median of -6.4 days from the observed peak, ranging individually from -9.8 to -5.3 days (S2 Table and S1 Fig). This is consistent with the findings in a human challenge study [26] where the viral load peak in the nose occurred 6.2 days after infection and ranged between 3 and 9 days. We note that the model predicted time to peak is not necessarily the same as the time to the observed peak viral load in the data. In some individuals the model predicts an earlier peak viral load compared to the observed peak viral load, e.g., the predicted viral load of individual 2349 peaks 4 days before the measured one (S1 Fig). In fact, the model predicts a time to peak of 5.7 days (Fig 3), i.e., on average 0.7 days before the observed peak, due to a rapid increase in viral load early post infection in some individuals. The model also predicts another 9.1 days to clear the infection (from peak viral load to the limit of detection) with a predicted infection duration of around 15 days.

**Figure 3:**
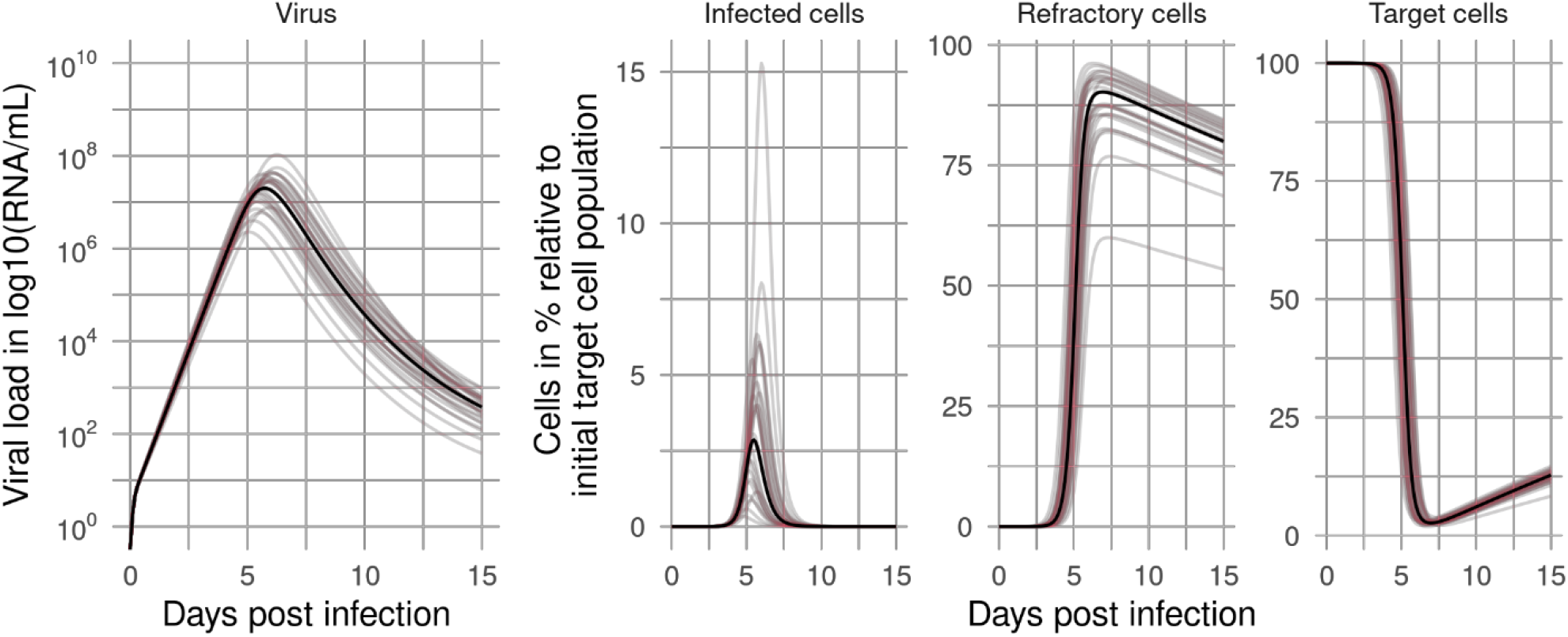
Virus and cell dynamics. Dynamics of virus, infected cells, refractory cells, and target cells (population = black line, individual = colored lines) throughout the course of infection predicted by the RCM using the best-fit population parameter estimates.

In combination with the eclipse phase duration, 1/*K*, the average lifespan of infected cells is 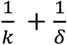. Using the estimated values of *K* and *δ*, we find the average lifespan of an infected cell is 0.64 days or 15 h. The within-host reproductive number *R*_0_ was on average 5 (Table 1).

**Table 1:** Parameters in the RCM viral dynamic model and their estimated population values. Values marked with * were fixed.

| Parameter | Description | RCM<br>Population estimate | Unit |
| --- | --- | --- | --- |
|  |  | [95% CI] |  |
| $T(t_{inf})$ | Initial target cell population | $8 \cdot 10^7^*$ | cells |
| $E(t_{inf})$ | Initial number of infected cells in the eclipse phase population | $1^*$ | cells |
| $t_{inf}$ | Infection time | -6.4<br>[-6.7, 6.1] | days |
| $\beta$ | Cell infection rate | $1.07 \cdot 10^{-8}$<br>[$9.12 \cdot 10^{-9}$, $1.26 \cdot 10^{-8}$] | mL/RNA<br>copies/day |
| $k$ | Transition rate out of the eclipse phase | $4^*$ | 1/day |
| $\pi$ | Virus production rate | 151<br>[131, 174] | RNA<br>copies/mL/day |
| $\delta$ | Death rate of infected cells | 2.58<br>[2.52, 2.64] | 1/day |
| $c$ | Virus clearance rate | $10^*$ | 1/day |
| $\varphi$ | Target to refractory cell conversion rate constant | $1.82 \cdot 10^{-6}$<br>[$1.25 \cdot 10^{-6}$, $2.69 \cdot 10^{-6}$] | 1/cell/day |
| $\rho$ | Refractory to target cell conversion rate | 0.016<br>[0.014, 0.019] | 1/day |
| $R_0$ | Basic reproductive number | 5.0 | |
| RMSE | Root mean squared error<br>Sum over all individuals: | 25.5 |  |
|  | Averaged over individuals: | 1.02 |  |
| -LL | negative log likelihood: | 921.3 |  |
| BICc | corrected Bayesian Information Criterion; | 980.8 |  |

The infected cell population peaks around 5.5 dpi (Fig 3). Target cells start to decline 3 to 4 dpi until depletion with less than 3% target cells left around 7 dpi when refractory cells reach their maximum of 81% of the initial susceptible target cells (Fig 3).

To further explore these fits, we performed a correlation analysis of the population parameters obtained from fits with a negative log-likelihood (-LL) in the range of 2 units from the best fit [that is, min(-LL) to min(-LL) + 2]. We found more than 50 fits with a -LL in the defined range with several model parameters significantly correlated (Fig 4). For example, the cell infection rate constant (*β*) and the virus production rate (*π*) are negatively correlated, as has been seen before [27]. The transition rate of susceptible cells into refractory cells (*φ*) is positively correlated with *π* but negatively correlated with *β*. Thus, the faster the estimated rate cells transition into the refractory state, the lower the estimated cell infection rate and the higher the estimated virus production rate. Furthermore, the transition rate of refractory cells back into susceptible cells (*ρ*) is positively correlated with the loss rate of infected cells (*δ*). Note that when we included these correlations in the model fitting, there was an increase in the BICc (991 with correlations compared to 981 without), and thus we did not include the correlations in further analyses.

**Figure 4:**
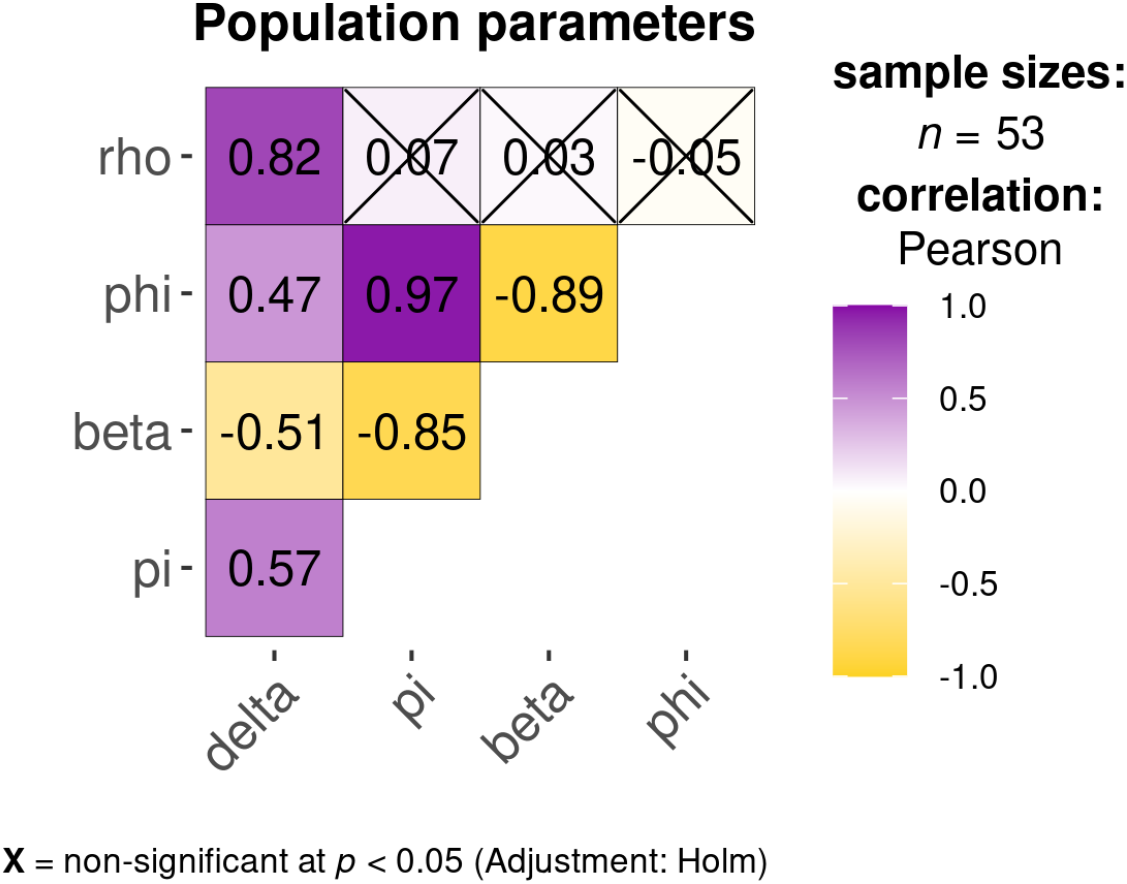
Correlation of population parameters in the refractory cell model. The sample size gives the number of fits that fit the model equally well in the range of min (-LL) to min(-LL)+2. Correlations that are crossed are non-significant (p-value > 0.05). The plot has been generated with ggstatsplot [doi:10.21105/joss.03167]

### Missing data in the exponential growth phase

In most human studies, data is not collected starting at the time of infection, but rather starting at the time of or later than the onset of symptoms [28]. To understand the effect of not having early data, we constructed different data sets, with varying numbers of viral load measurements during the viral up-slope, to study the robustness of estimated model parameters to missing data by comparison with the results obtained by fitting the full data set presented above. Based on the estimate of *t*_*inf*_ for each individual, data sets were constructed starting i) 3 days post infection (dpi), yielding on average 2.1 ± 1.6 pre-peak measurements, ii) 5 dpi, yielding on average 0.7 ± 0.9 pre-peak measurements, and iii) 7 dpi yielding on average 0.1 ± 0.6 pre-peak measurements. In the 3-, 5-, and 7-dpi data sets, pre-peak viral load measurements were available from 21, 12, and 1 individual, respectively. Note that the 7-dpi data set starts very near the peak viral load (±1 day), and only one individual has pre-peak data, 17 individuals lose the peak viral load, and 5 individuals have missing data after peak viral load (viral down-slope). We further studied the robustness of parameter estimates using only the peak and post-peak measurements, as a proxy for data collection around symptom onset and, consequently, the most common data set obtained in clinical practice.

An important issue with fitting acute infection data is that typically we do not know the time of infection, and thus don’t know the times relative to infection when data was collected. To study this issue, we considered three different scenarios for each of the artificial data sets created above. First, we assumed that we do not know the infection time and estimate it (*t*_*inf*_ = *est*) from the data as we did above, but now using our reduced data sets. Here the first measurement is assigned time 0 and we estimate infection before that. In the second case, we assume we know the time of infection, as estimated from the full data set, and, thus, *t*_*inf*_ = 0 is the actual infection time. We simply fit the model to the various data sets and estimate model parameters (with *t*_*inf*_ = 0). Lastly, when data in the exponential growth phase is missing, it is common practice to set the time of infection from the literature [25,29–32], e.g., by assuming viral load peaks at the estimated median time of symptom onset, i.e., 5 dpi for SARS-CoV-2 [13,14]. Thus, for this third case, we only use the post-peak data set and assume the time of the observed peak viral load is day 5.

We fitted the model in turn to these different data sets (and assumptions of *t*_*inf*_) and then use all available data points above the LOD to calculate the RMSE, as we did for the full model fit. Therefore, the lower the RMSE, the more the predictions of a model fitted to a subset of the data agree with the full viral load course. Fitting the model to these modified data sets, we found, as expected, that the more data available in the exponential growth phase (pre-peak), the lower the RMSE and the more reliable the estimated course of infection (S2 Fig). However, since infection times and the number of days missing in the exponential growth phase are often unknown, we averaged the RMSEs over the fits obtained for the 3, 5, 7 dpi, and post-peak data sets to get an idea of how much data we need for the model to perform well under different assumptions for the time of infection.

Knowing the time of infection (*t*_*inf*_ = 0) generally results in lower RMSEs than when estimating *t*_*inf*_ (Fig 5, black squares). Furthermore, if we assume that the viral load peaks 5 dpi and fix this (Fig 5, blue dots), we obtain lower RMSEs compared to re-estimating infection times for each individual (Fig 5, orange triangles). The TCLM yields similar results, but interestingly, in many cases that model also yields slightly lower average RMSEs than the RCM (S1 Text).

**Figure 5:**
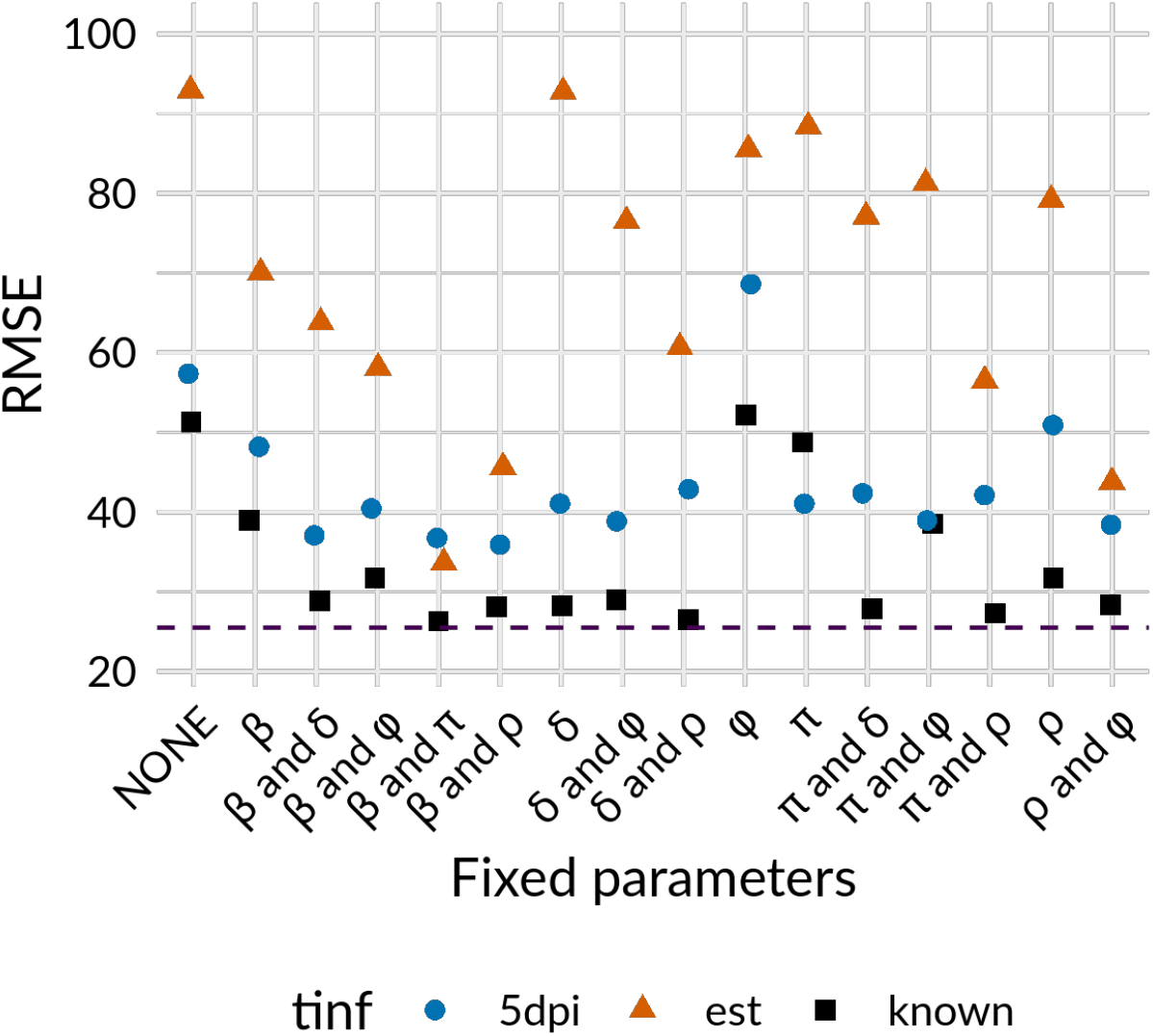
RMSEs for the RCM and three infection time cases. Infection times (t_inf_) are re-estimated (triangle), infection times (t_inf_) known and set to zero (square), or infection times are set to zero and the VL peaks 5 dpi (circle). RMSEs are averaged over the different data collection scenarios (3-, 5-, 7-dpi, and post peak). For each case, all model parameters are re-estimated (NONE on x-axis), or model parameters were fixed to the values estimated from the full course of infection data set (see Table 1). The dashed line represents the RMSE calculated from the best model fit using the full course of infection. (The corresponding plot for individual data collection scenarios are shown in Fig S2 and population parameters estimated for the different scenarios can be found in Fig S3).

Fixing one or more model parameters is common practice to reduce uncertainty in data fitting [33]. Therefore, we were interested in how many model parameters in addition to *c* and *KK* had to be fixed to describe well the full course of infection with the different data sets. The parameters *cc* and *KK* are typically fixed as they refer to processes that take place on timescale of minutes to hours and for which data on these timescales is unavailable [15,24,25,34–37]. Thus, we systematically fixed every single or possible pair of remaining model parameters to the population value estimated from the entire data set representing the entire course of infection (Table 1). Adding knowledge to the model fitting by fixing one or two additional model parameters improved the RMSEs. Knowing *t*_*inf*_ yielded the lowest RMSE, followed by assuming the viral load peaks 5 dpi, where both outperformed re-estimating *t*_*inf*_. Especially by fixing the cell infection rate *β* and the virus production rate *π*, we observed overall the lowest RMSEs, which were close to those calculated from the whole course of infection data set (Fig 5). Both are crucial parameters of the exponential growth phase. However, fixing only one of those model parameters (*β* or *π*) led to conflicting results (Fig. 5). If infection times are not known and, thus, must be estimated, fixing only *β* yielded lower RMSEs than fixing only *π*. However, if we assume the viral load peaks 5 dpi, fixing *π* yielded lower RMSEs than fixing *β* (Fig 5). Additionally, if infection times are re-estimated or if we assume the viral load peaks 5 dpi and fix *π*, the TCLM yielded lower RMSEs than the RCM (S1 Text).

### How reliable are estimated infection times and other model parameters?

To evaluate the robustness of model parameter estimates, we estimated them from the modified data sets and compared them to the model parameters estimated from the full course of infection with and without fixing model parameters (beyond *c* and *k*).

The estimated *δ* was close to the value estimated from the full course of infection if *ρ* was fixed due to their correlation (Fig 6A and S3 Fig). Furthermore, only by fixing *β* and *π*, we were able to accurately estimate the time of infection reliably and almost exactly (Fig 6B). Estimating *β* was most reliable if we assume the viral load peaks 5 dpi (Fig 6C). Again, *π* was mostly over or underestimated when fixing one or two model parameters. However, not fixing any parameters led to the most reliable estimate of *π* for all three studied cases (Fig 6D). For both innate immune response model parameters *φ* and *ρ*, fixing *β*and *π* or *β* and *δ* performed best for all data collection scenarios (Fig 6E and 6F).

**Figure 6:**
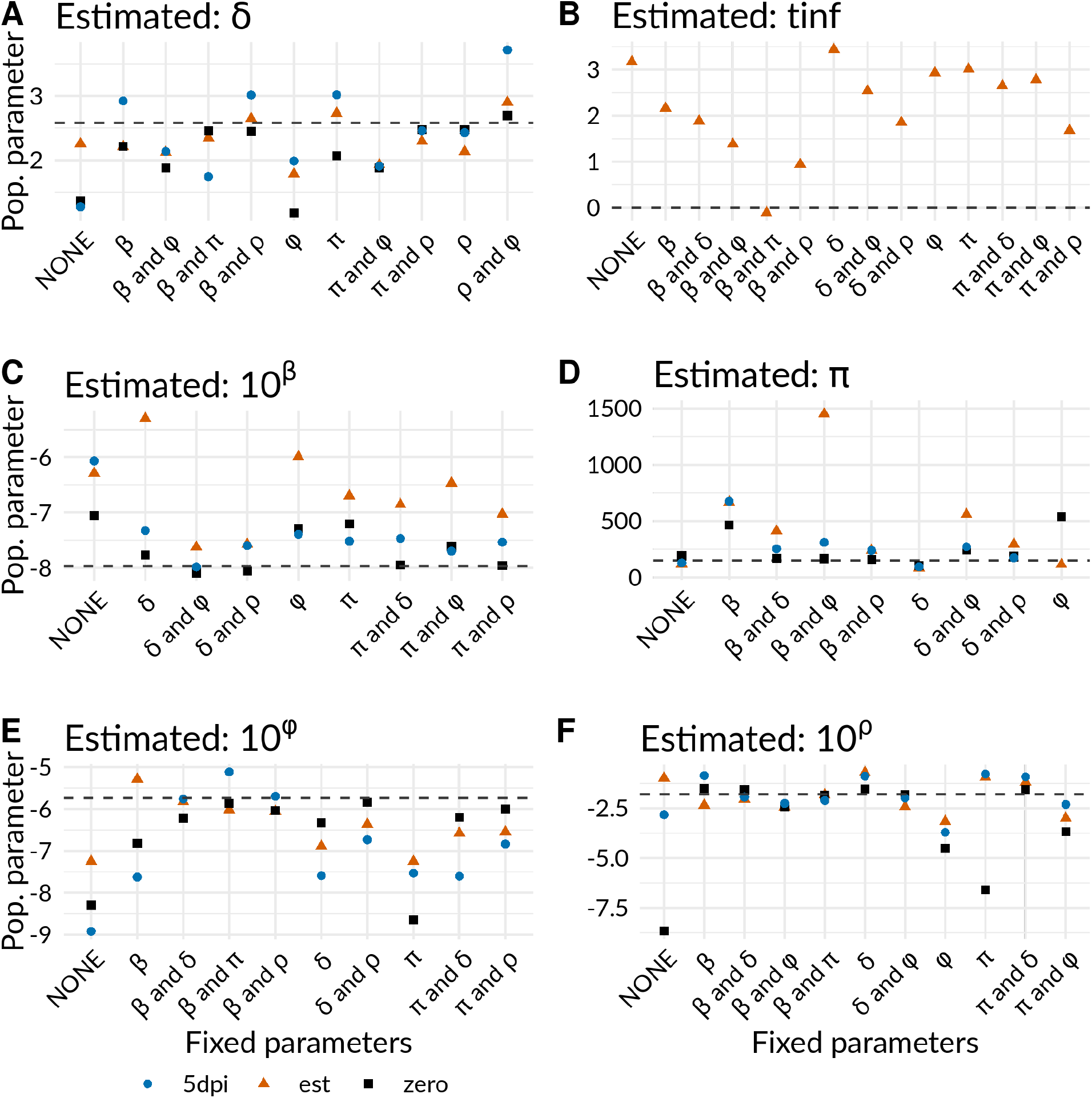
Estimated population parameters averaged over the four data collection scenario using the RCM. The dotted line represents the population parameter estimated from the “full course of infection” data set (Population parameters estimated from every data collection scenario can be found in Fig S3).

### Effect of choosing different combinations of model parameters

Lastly, we were interested in the model performance by choosing different combinations of the cell infection rate (*β*) and the virus production rate (*π*), beyond what we estimated in Table 1. For that, we calculated the RMSEs of model fits where the cell infection rate (*β*) and the virus production rate (*π*) where fixed and the remaining model parameters were estimated. For these analyses, we used the post-peak data set and assumed 5 days to reach peak viral load. As shown in Fig 7, *β* and *π* correlate inversely and, because of this, we found that different combinations of *β* and *π* led to equally good fits, represented as dark blue tiles and, thus, low RMSEs (Fig 7).

**Figure 7:**
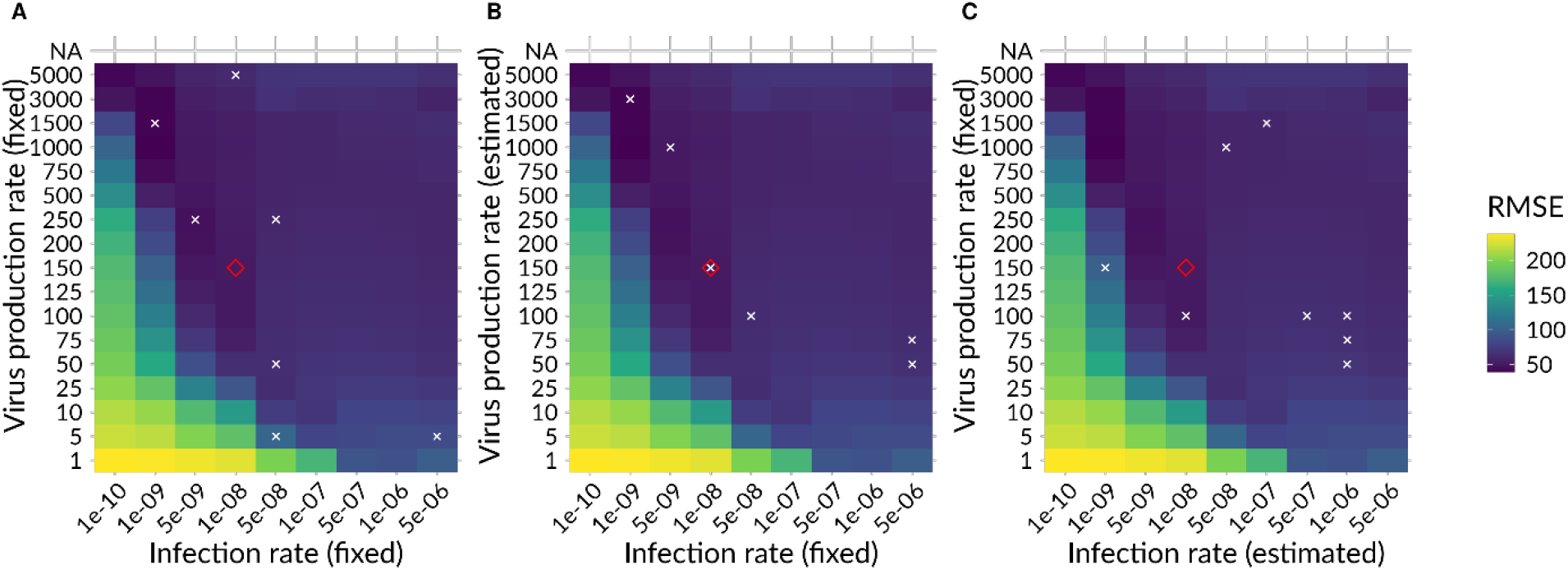
Heatmap RMSEs calculated with the RCM and different combinations of population parameters. A) RMSEs for different combinations of literature values. B) RMSEs for fixed infection rates from literature and estimated virus production rates. C) RMSEs for fixed virus production rates from literature and estimated infection rates. ◊ = the population parameters we estimated, x = values for β and/or π found in literature. Parameter values can be found in Table S3.

Furthermore, since *β* and *π* are often unknown and challenging to measure experimentally, we tested our model performance by fixing *β* and/or *π* to different values from the literature [1,12,29,38–42] (S3 Table, Fig 7). Compared to our estimate of *β* = 1 · 10^−8^ mL/RNA copies/day, the infection rate values found in literature were mostly in the range between 1 · 10^−9^ to 5 · 10^−8^. These values and their corresponding virus production rates, which ranged on average between 50 and 1500 (our estimate *π* = 151 RNA copies/mL/day), led to equally good fits (Fig 7A). However, we observed overall lower RMSEs, if fixing *β* and estimating *π* (Fig 7B) instead of fixing *π* and estimating *β* (Fig 7C). Thus, using infection rate values from the literature represents a good strategy to deal with missing data in the exponential growth phase and missing information about infection times.

## Discussion

Reliably estimating parameter values in viral dynamic models with missing data is challenging. Especially in acute infections, where individuals generally only become aware of being infected when symptoms develop. Thus, information about the time of infection and viral load measurements prior to symptom onset is often not available. In the present study, we analyzed the reliability of estimated viral dynamics model parameters in the absence of variable amounts of data in the exponential growth phase. We found that viral infection and production rates are key parameters in determining the exponential growth rate. Especially with a lack of early data, the time to peak viral load was often underestimated. However, fixing the time of infection based on epidemiological studies represented a good alternative to estimating infection times and resulted in good model fits.

### Viral dynamics of the entire course of infection

The RCM describes the frequent viral load measurements of the 25 studied individuals well. Most estimated model parameters agreed with our previous work, except for the transition rate turning refractory cells back into susceptible cells, which we now estimate almost 3-fold higher [1]. Interestingly, we estimated that only 3% of the total cells were infected at the peak and 6% cumulative from infection to peak viral load. However, at the peak viral load most cells were in a refractory state (81%) and 12% of cells remaining susceptible to infection. Turning target cells into cells refractory to viral infection by establishing an antiviral state in uninfected cells may be a critical host defense mechanism early on in fighting a viral infection. However, as far as we know experimental measurements of the fraction of cells in an antiviral state during SARS-CoV-2 infection are not available and thus limit our ability to compare these predictions to data.

### The effect of missing data in the exponential growth phase

With missing data in the exponential growth phase, infection times are underestimated by 2 to 3 days, resulting in very fast estimated initial growth rates. However, we improved the infection time estimates by adding knowledge to the model. Cell infection and virus production rates are crucial parameters for describing the exponential growth phase. Fixing both model parameters to our population values led to reliable infection time estimates similar to those estimated from the entire course of infection data set.

We were further interested in the reliability of other model parameter estimates. By fitting the RCM to different data collection scenarios, we found that knowing the infection times led to the lowest RMSEs. However, a low RMSE did not guarantee the correct estimation of population parameters due to the correlations in the model structure such as the correlation between the cell infection rate (*β*) and the virus-production rate (*π*). Furthermore, since infection times are often unknown, estimating infection times or having an idea about them from epidemiological studies and fixing them are more realistic but led to higher RMSEs. Estimated infection times were underestimated by up to 3 days, while fixing *β* and *π* led to the most robust infection time-estimates. Nevertheless, assuming a time to peak viral load of 5 days (for SARS-CoV-2) represented a good alternative to estimating infection times and estimated population parameters close to those estimated from the full course of infection.

Interestingly, whether infection times are known, estimated or assumed, the loss rate of infected cells represented the most robust model parameter, with more consistent estimation, due to frequent viral load measurements after symptom onset and thus after peak viral load.

### Estimating the exponential growth phase parameters: What if β and π are unknown?

The cell infection rate *β* and virus production rate *π* are crucial parameters of the exponential viral growth phase. Fixing both model parameters may lead to reliable predictions of infection times. However, both model parameters are often unknown and challenging to measure experimentally.

If infection times are unknown, assuming 5 days from the time of infection to peak viral load led to the most reliable estimates of *β*. Nevertheless, estimates of *π* showed more variability, which may be due to lower sensitivity. It has been further shown that the initial target cell population *T*(*t*_*inf*_) also correlates with the virus production rate *π* and only their product [*T*(*t*_*inf*_)· *π*] is identifiable [43,44].

However, estimates for cell infection and virus production rates from other modeling studies [1,12,29,38– 42] fit our data equally well, such as *β* = 10^−8^ mL/RNA copies/day and *π* = 150 to 200 RNA copies/mL/ day or *β* = 10^−9^ mL/RNA copies/day and *π* = 1000 to 1500 RNA copies/mL/ day. Consequently, with missing data in the exponential growth phase taking cell infection and virus production rates from the literature may allow robust predictions of the exponential growth phase.

### Limitations and outlook

Our analysis was based on models of acute infection that have been used for a variety of viruses including West Nile virus [45], respiratory syncytial virus [46], influenza [15,27,34,47,48], and SARS-CoV-2 [1,12,17,25,39]. However, here we only analyzed data for SARS-CoV-2 infection due to the availability of a rich dataset. Also, we selected our data from a unique cohort that included primarily male, young, healthy, and physically active athletes. However, vendors and staff were also regularly tested and part of the data set. Even though, the cohort may not be representative of the total population of infected individuals, no difference in viral load of different age or demographic groups has been reported [12]. Thus, the conclusions made in the presented analysis will not be affected by the bias in the cohort we used. Instead, our conclusions inform about the reliability of model parameter estimates in general and may be particularly beneficial for respiratory infections.

Furthermore, future epidemics and pandemics are inevitable, and our results may be useful in terms of guiding data collection and in using that data to best estimate viral dynamic parameters such as the death rate of infected cells, which can inform us about viral pathogenesis. Moreover, we emphasize that only with the most informative data sets, i.e., frequent measurements throughout the course of infection, can we accurately infer the infection kinetics and the infectious period of an individual if a novel respiratory virus emerges in the future.

In summary, the current study provides new insights into viral dynamic modeling in the absence of frequent viral load measurements. We evaluated the reliability of estimated model parameters and found that cell infection and virus production rates are key parameters of the exponential viral growth phase. Furthermore, missing data before the viral load peaks leads to underestimates of the time to peak viral load and to unreliable estimated model parameters. However, fixing infection times from epidemiological studies, and model parameters of the exponential growth rate (*β* and *π*) represented a good alternative to estimating infection times and led to good model fits and model parameters estimates.

---

***S1 Table: Sensitivity of the fit to the initial number of infected cells E***(**t**_*inf*_). *Highlighted in orange is the best fit*. *[-LL = negative log likelihood*, *BICc = corrected Bayesian Information Criterion*, *TCLM = Target cell limited model*, *RCM = Refractory cell model]*

***S2 Table: Individual parameters in the RCM and their estimated values***.

***S3 Table: Estimated beta and pi parameter values from literature***. *TCLM = Target cell limited model*, *RCM = Refractory cell model*

***S1 Fig: Best model fit with peak viral load at t = 0***. *The best model fit to viral load measurements of 25 selected individuals with t* = 0 *corresponds to the measured peak viral load*. *Filled circles are measurement points*, *and non-filled circles are censored and below the detection limit (dotted grey line)*.

***S2 Fig: RMSEs for RCM and data collected 3***, ***5***, ***7***, ***dpi***, ***or post-peak***. *RMSEs for RCM and the different data collection scenarios and A) infection times (t*_*inf*_ *) are re-estimated or B) infection times (t*_*inf*_*) are known and set to zero*. *For each data set*, *all model parameters are re-estimated (NONE on x-axis)*, *or model parameters were fixed to the values estimated from the full course of infection data set (see Table 1)*. *The dashed line represents the RMSE calculated from the best model fit using the full course of infection*.

***S3 Fig: Estimated population parameters of the RCM and data collected 3***, ***5***, ***7***, ***dpi***, ***or post-peak***. *The dotted line represents the population parameter estimated from the full course of infection data set (Y axis are estimated population values)*.

***S1 Text: The target cell limited model***.

## References

1. Ke R, Zitzmann C, Ho DD, Ribeiro RM, Perelson AS. In vivo kinetics of SARS-CoV-2 infection and its relationship with a person’s infectiousness. Proc Natl Acad Sci U S A. 2021;118. doi:10.1073/PNAS.2111477118/-/DCSUPPLEMENTAL

2. Marc A, Kerioui M, Blanquart F, Bertrand J, Mitjà O, Corbacho-Monné M, et al. Quantifying the relationship between sars-cov-2 viral load and infectiousness. Elife. 2021;10. doi:10.7554/ELIFE.69302

3. Wölfel R, Corman VM, Guggemos W, Seilmaier M, Zange S, Müller MA, et al. Virological assessment of hospitalized patients with COVID-2019. Nature 2020 581:7809. 2020;581: 465–469. doi:10.1038/s41586-020-2196-x

4. Carrat F, Vergu E, Ferguson NM, Lemaitre M, Cauchemez S, Leach S, et al. Time Lines of Infection and Disease in Human Influenza: A Review of Volunteer Challenge Studies. Am J Epidemiol. 2008;167: 775–785. doi:10.1093/AJE/KWM375

5. Tsang TK, Cowling BJ, Fang VJ, Chan KH, Ip DKM, Leung GM, et al. Influenza A Virus Shedding and Infectivity in Households. J Infect Dis. 2015;212: 1420–1428. doi:10.1093/INFDIS/JIV225

6. Asher J, Lemenuel-Diot Id A, Clay Id M, Durham DP, Mier-Y-Teran-Romeroid L, Arguello CJ, et al. Novel modelling approaches to predict the role of antivirals in reducing influenza transmission. PLoS Comput Biol. 2023;19:p e1010797. doi:10.1371/JOURNAL.PCBI.1010797

7. Eisinger RW, Dieffenbach CW, Fauci AS. HIV Viral Load and Transmissibility of HIV Infection: Undetectable Equals Untransmittable. JAMA. 2019;321: 451–452. doi:10.1001/JAMA.2018.21167

8. Fraser C, Lythgoe K, Leventhal GE, Shirreff G, Hollingsworth TD, Alizon S, et al. Virulence and pathogenesis of HIV-1 infection: an evolutionary perspective. Science. 2014;343. doi:10.1126/SCIENCE.1243727

9. Wilson DP, Law MG, Grulich AE, Cooper DA, Kaldor JM. Relation between HIV viral load and infectiousness: a model-based analysis. The Lancet. 2008;372: 314–320. doi:10.1016/S0140-6736(08)61115-0

10. Kissler SM, Fauver JR, Mack C, Olesen SW, Tai C, Kalinich CC, et al. SARS-CoV-2 viral dynamics in acute infections. medRxiv. 2020; 2020.10.21.20217042. doi:10.1101/2020.10.21.20217042

11. Kissler SM, Fauver JR, Mack C, Tai CG, Breban MI, Watkins AE, et al. Viral dynamics of SARS-CoV-2 variants in vaccinated and unvaccinated persons. New England Journal of Medicine. 2021;385: 2489–2491. doi:10.1056/nejmc2102507

12. Ke R, Martinez PP, Smith RL, Gibson LL, Mirza A, Conte M, et al. Daily longitudinal sampling of SARS-CoV-2 infection reveals substantial heterogeneity in infectiousness. Nature Microbiology 2022 7:5. 2022;7: 640–652. doi:10.1038/s41564-022-01105-z

13. Sanche S, Lin YT, Xu C, Romero-Severson E, Hengartner N, Ke R. High Contagiousness and Rapid Spread of Severe Acute Respiratory Syndrome Coronavirus 2. Emerg Infect Dis. 2020;26: 1470–1477. doi:10.3201/eid2607.200282

14. Lauer SA, Grantz KH, Bi Q, Jones FK, Zheng Q, Meredith HR, et al. The Incubation Period of Coronavirus Disease 2019 (COVID-19) From Publicly Reported Confirmed Cases: Estimation and Application. Ann Intern Med. 2020;172: 577–582. doi:10.7326/M20-0504

15. Baccam P, Beauchemin C, Macken CA, Hayden FG, Perelson AS. Kinetics of influenza A virus infection in humans. J Virol. 2006;80: 7590–9. doi:10.1128/JVI.01623-05

16. Best K, Guedj J, Madelain V, de Lamballerie X, Lim S-Y, Osuna CE, et al. Zika plasma viral dynamics in nonhuman primates provides insights into early infection and antiviral strategies. Proc Natl Acad Sci U S A. 2017;114: 8847–8852. doi:10.1073/pnas.1704011114

17. Perelson AS, Ke R. Mechanistic modelling of SARS-CoV-2 and other infectious diseases and the effects of therapeutics. Clin Pharmacol Ther. 2021. doi:10.1002/cpt.2160

18. Zitzmann C, Kaderali L. Mathematical analysis of viral replication dynamics and antiviral treatment strategies: From basic models to age-based multi-scale modeling. Front Microbiol. Frontiers; 2018. p. 1546. doi:10.3389/fmicb.2018.01546

19. Banerjee S, Guedj J, Ribeiro RM, Moses M, Perelson AS. Estimating biologically relevant parameters under uncertainty for experimental within-host murine West Nile virus infection. J R Soc Interface. 2016;13. doi:10.1098/RSIF.2016.0130

20. Samuel CE. Antiviral actions of interferons. Clinical Microbiology Reviews. American Society for Microbiology (ASM); 2001. pp. 778–809. doi:10.1128/CMR.14.4.778-809.2001

21. Levy DE, GarcÍa-Sastre A. The virus battles: IFN induction of the antiviral state and mechanisms of viral evasion. Cytokine Growth Factor Rev. 2001;12: 143–156. doi:10.1016/S1359-6101(00)00027-7

22. GarcÍa-Sastre A, Biron CA. Type 1 Interferons and the Virus-Host Relationship: A Lesson in Détente. Science (1979). 2006;312: 879–882. doi:10.1126/SCIENCE.1125676

23. Ke R, Zitzmann C, Ribeiro RM, Perelson AS. Kinetics of SARS-CoV-2 infection in the human upper and lower respiratory tracts and their relationship with infectiousness. medRxiv. medRxiv; 2020. p. 2020.09.25.20201772. doi:10.1101/2020.09.25.20201772

24. Smith AP, Moquin DJ, Bernhauerova V, Smith AM. Influenza virus infection model with density dependence supports biphasic viral decay. Front Microbiol. 2018;9. doi:10.3389/FMICB.2018.01554

25. Gonçalves A, Bertrand J, Ke R, Comets E, Lamballerie X, Malvy D, et al. Timing of Antiviral Treatment Initiation is Critical to Reduce SARS-CoV-2 Viral Load. CPT Pharmacometrics Syst Pharmacol. 2020;9: 509–514. doi:10.1002/psp4.12543

26. Killingley B, Mann A✉. Safety, tolerability and viral kinetics during SARS-CoV-2 human challenge in young adults. [cited 10 Aug 2023]. doi:10.1038/s41591-022-01780-9

27. Smith AM. Host-pathogen kinetics during influenza infection and coinfection: insights from predictive modeling. Immunol Rev. 2018;285: 97–112. doi:10.1111/IMR.12692

28. Challenger JD, Foo CY, Wu Y, Yan AWC, Marjaneh MM, Liew F, et al. Modelling upper respiratory viral load dynamics of SARS-CoV-2. BMC Med. 2022;20. doi:10.1186/S12916-021-02220-0

29. Hernandez-Vargas EA, Velasco-Hernandez JX. In-host Mathematical Modelling of COVID-19 in Humans. Annu Rev Control. 2020;50: 448–456. doi:10.1016/J.ARCONTROL.2020.09.006

30. Goyal A, Cardozo-Ojeda EF, Schiffer JT. Potency and timing of antiviral therapy as determinants of duration of SARS-CoV-2 shedding and intensity of inflammatory response. Sci Adv. 2020;6. doi:10.1126/SCIADV.ABC7112/SUPPL_FILE/ABC7112_SM.PDF

31. Sadria M, Layton AT. Modeling within-Host SARS-CoV-2 Infection Dynamics and Potential Treatments. Viruses 2021, Vol 13, Page 1141. 2021;13:p 1141. doi:10.3390/V13061141

32. Gonçalves A, Bertrand J, Ke R, Comets E, de Lamballerie X, Malvy D, et al. Timing of Antiviral Treatment Initiation is Critical to Reduce SARS-CoV-2 Viral Load. CPT Pharmacometrics Syst Pharmacol. 2020;9:p 509. doi:10.1002/PSP4.12543

33. Maiwald T, Hass H, Steiert B, Vanlier J, Engesser R, Raue A, et al. Driving the Model to Its Limit: Profile Likelihood Based Model Reduction. 2016 [cited 3 Feb 2023]. doi:10.1371/journal.pone.0162366

34. Pawelek KA, Huynh GT, Quinlivan M, Cullinane A, Rong L, Perelson AS. Modeling Within-Host Dynamics of Influenza Virus Infection Including Immune Responses.Antia R, editor. PLoS Comput Biol. 2012;8:p e1002588. doi:10.1371/journal.pcbi.1002588

35. Néant N, Lingas G, Le Hingrat Q, Ghosn J, Engelmann I, Lepiller Q, et al. Modeling SARS-CoV-2 viral kinetics and association with mortality in hospitalized patients from the French COVID cohort. Proceedings of the National Academy of Sciences. 2021;118:p e2017962118. doi:10.1073/pnas.2017962118

36. Hou YJ, Okuda K, Edwards CE, Martinez DR, Asakura T, Dinnon KH, et al. SARS-CoV-2 Reverse Genetics Reveals a Variable Infection Gradient in the Respiratory Tract. Cell. 2020;182:p 429. doi:10.1016/J.CELL.2020.05.042

37. Ogando NS, Dalebout TJ, Zevenhoven-Dobbe JC, Limpens RWAL, van der Meer Y, Caly L, et al. SARS-coronavirus-2 replication in Vero E6 cells: replication kinetics, rapid adaptation and cytopathology. J Gen Virol. 2020;101:p 925. doi:10.1099/JGV.0.001453

38. Goyal A, Reeves DB, Fabian Cardozo-Ojeda E, Schiffer JT, Mayer BT. Viral load and contact heterogeneity predict sars-cov-2 transmission and super-spreading events. Elife. 2021;10: 1–63. doi:10.7554/ELIFE.63537

39. Kim KS, Ejima K, Iwanami S, Fujita Y, Ohashi H, Koizumi Y, et al. A quantitative model used to compare within-host SARS-CoV-2, MERS-CoV, and SARS-CoV dynamics provides insights into the pathogenesis and treatment of SARS-CoV-2. PLoS Biol. 2021;19:p e3001128. doi:10.1371/journal.pbio.3001128

40. Padmanabhan P, Dixit N. Modeling Suggests a Mechanism of Synergy Between Hepatitis C Virus Entry Inhibitors and Drugs of Other Classes. CPT Pharmacometrics Syst Pharmacol. 2015;4: 445–453. doi:10.1002/psp4.12005

41. Perelson AS, Ribeiro RM, Phan T. An explanation for SARS-CoV-2 rebound after Paxlovid treatment. medRxiv. 2023; 2023.05.30.23290747. doi:10.1101/2023.05.30.23290747

42. Ejima K, Kim KS, Ludema C, Bento AI, Iwanami S, Fujita Y, et al. Estimation of the incubation period of COVID-19 using viral load data. Epidemics. 2021;35:p 100454. doi:10.1016/J.EPIDEM.2021.100454

43. Stafford MA, Corey L, Cao Y, Daar ES, Ho DD, Perelson AS. Modeling plasma virus concentration during primary HIV infection. J Theor Biol. 2000;203: 285–301. doi:10.1006/jtbi.2000.1076

44. Miao H, Xia X, Perelson AS, Wu H. On Identifiability of Nonlinear ODE Models and Applications in Viral Dynamics. https://doi.org/101137/090757009. 2011;53:p 3–39. doi:10.1137/090757009

45. Banerjee S, Guedj J, Ribeiro RM, Moses M, Perelson AS. Estimating biologically relevant parameters under uncertainty for experimental within-host murine West Nile virus infection. J R Soc Interface. 2016;13. doi:10.1098/RSIF.2016.0130

46. Patel K, Kirkpatrick CM, Nieforth KA, Chanda S, Zhang Q, McClure M, et al. Respiratory syncytial virus-A dynamics and the effects of lumicitabine, a nucleoside viral replication inhibitor, in experimentally infected humans. Journal of Antimicrobial Chemotherapy. 2019;74: 442–452. doi:10.1093/JAC/DKY415

47. Hadjichrysanthou C, Cauët E, Lawrence E, Vegvari C, de Wolf F, Anderson RM, et al. Understanding the within-host dynamics of influenza A virus: from theory to clinical implications. Journal of the Royal Society, Interface /the Royal Society. 2016;13: 499–522. doi:10.1098/rsif.2016.0289

48. Smith AM, Perelson AS. Influenza A virus infection kinetics: Quantitative data and models. Wiley Interdiscip Rev Syst Biol Med. 2011;3: 429–445. doi:10.1002/wsbm.129

